# Butyrate decreases *Campylobacter jejuni* motility and attachment partially through influence on LysR expression

**DOI:** 10.1101/2022.09.29.510231

**Authors:** Nereus W. Gunther, Alberto Nunez, Lori Bagi, Aisha Abdul-Wakeel, Amy Ream, Yanhong Liu, Gaylen Uhlich

## Abstract

The food pathogen *Campylobacter jejuni* both colonizes the lower intestines of poultry and infects the lower intestines of humans. The lower intestines of both poultry and humans are also home to a wide range of commensal organisms which compete with an organism like *C. jejuni* for space and resources. The commensal organisms are believed to protect humans against infection by pathogens of the digestive tract like *C. jejuni*. The short chain fatty acid (SCFA) butyrate is a metabolite commonly produced by commensal organisms within both the poultry and human digestive tract. We investigated the affect that physiologically relevant concentrations of butyrate have on *C. jejnui*. Butyrate at concentrations of 5 and 20 mM negatively impacted *C. jejuni* motility and attachment. These two traits are believed important for *C. jejuni’s* ability to infect the lower intestines of humans. Additionally, 20 mM butyrate concentrations were observed to influence the expression of a range of different *Campylobacter* proteins. Constitutive expression of one of these proteins, LysR, within a *C. jejuni* strain partially lessened the negative influence butyrate had on the bacteria’s motility.

**Importance:** Research studies on the short chain fatty acid, butyrate, have produced evidence for it both negatively and positively impacting *Campylobacter jejuni’s* colonization of poultry or infection of the human intestinal tract. There is significant value in clarifying if butyrate has potential as an intervention agent capable of interfering in the virulence process of *C. jejuni*. The results presented, show that butyrate, at physiological levels, negatively impacts *C. jejuni* motility and attachment, two important factors in the virulence process. Additionally, a proteomic analysis of butyrate’s influence on *C. jejuni*, identified that the expression of the transcriptional factor, LysR, was repressed in the presence of butyrate. Overexpression of the *lysR* gene restored some motility in the presence of butyrate suggesting this factor had a positive influence on the flagella necessary for *C. jejuni’s* movement.

## Introduction

Among cases of bacteria mediated foodborne gastrointestinal diseases of humans, *Campylobacter* stands out as causing the largest number of infections annually in the developed world (1, 2). The genus *Campylobacter* currently contains four species (*jejuni, coli, lari* and *upsaliensis*) known to regularly cause human disease, however among these four, *C. jejuni* is roughly responsible for 90% of the disease cases (2-4).

*Campylobacter jejuni* regularly colonizes the digestive tracts of chickens raised for human consumption, leading to this food product being contaminated with the bacteria during the slaughter process (5, 6). This results in poultry being the primary source for transmission of *C. jejuni* to humans through the consumption of undercooked poultry or by cross contamination of other food products (7, 8). The lower intestinal tracts of both poultry and humans are an environmental niche regularly exploited by *C. jejuni*. Additionally, chickens’ and humans’ lower intestinal tracts are occupied by a diverse population of commensal bacterial strains producing a host of metabolites (9-11). Among the bacterial produced metabolites are short chain fatty acids (SCFAs), which are found in high concentrations in the lower intestines of both chickens and humans, with the most prevalent forms being acetate, butyrate and propionate (12). Recent research studies have made a compelling case for *C. jejuni* utilizing the sensing of the SCFA, butyrate, to preferentially colonize the lower intestines of chickens or infect the lower intestines of humans (13, 14). The researcher identified a sensor in *C. jejuni* that detects the presence of butyrate and appears to up regulate genes required for successful colonization of hosts (13). However, previous studies looking at different compositions of chicken feed, identified feed compositions that increased SCFA concentrations that also led to a decrease in *C. jejuni* colonization of the chicken digestive tract (15). Additionally, another research group presented evidence for butyrate and other SCFAs providing protection against *C. jejuni* colonization of chickens and invasion of human cells (16, 17).

The response of two other food pathogens, *Escherichia coli* and *Salmonella*, to SCFAs has also been studied and the bacterial sensors for detecting SCFAs described (18, 19). Contrary to the previous suggested role for butyrate enhancing colonization by *C. jejuni*, butyrate’s interaction with *Salmonella* appears to reduce the virulence potential of that pathogen (20). However, the enterohaemorrhagic class of *E. coli* when exposed to butyrate begin to upregulate a range of known virulence factors (21). The contradictory research conclusions and different behavior among other food pathogens therefore places the role for butyrate’s influence on *C* .*jejuni* as needing additional research. A more recent research paper observed that butyrate increased the motility of *E. coli* by enhancing flagellar expression (22). Motility and flagella production are factors that are well accepted as essential for both colonization and infection by *C. jejuni* (23, 24). Therefore, an investigation of butyrate’s influence on *C. jejuni’s* motility should serve to add weight to one side or the other in a discussion of whether the presence of butyrate plays a positive or a negative role in *C. jejuni’s* colonization or infection of the lower intestines of chickens and humans.

## Materials and Methods

### Bacterial strains and plasmids

The *Campylobacter jejuni* strains and plasmids used in this study are listed in Table 1. All strains were stored in Brucella broth (Becton Dickinson, Sparks, MD) plus 15% glycerol frozen at -80°C until required for experiments. Strains were struck directly from freezer stocks onto Brucella agar (1.5%) plates. Kanamycin (15 μg/mL final concentration) was added to any Brucella agar plates for growing strains possessing a cloned *aph*(3) gene. All *C. jejuni* grown for this paper were incubated at 42°C in a microaerobic (5% O_2_, 10% CO_2_, 85% N_2)_ growth chamber (Concept-M, Baker Ruskinn, UK).

**Table 1:** Strains and plasmids utilized in research.

| Strain | Description | Source or Reference |
| --- | --- | --- |
| *C. jejuni* CDC279 | Clinical isolate CDC279(2014D-0279) | Centers for Disease Control (CDC) |
| *C. jejuni* CDC9511 | Clinical isolate CDC9511(2012D-9511) | Centers for Disease Control (CDC) |
| *C. jejuni* RM3194 | Clinical isolate | (31) |
| *C. jejuni* RM3194+<br>express | RM3194 with an empty *PcysM* expression element integrated into chromosome; $\text{Kn}^{\text{R}}$ | This study |
| *C. jejuni* RM3194+*lysR*-express | RM3194 with *PcysM-lysR* expression element integrated into chromosome; $\text{Kn}^{\text{R}}$ | This study |
| Plasmid | Description | Source |
| pBlueKan+cysM<sup>Pro</sup> | pBluescriptII KS+ with cloned *aph*(3), *cysM* promoter, and unique 3' *Sma*I and *Xho*I cloning sites; $\text{Kn}^{\text{R}}$; $\text{Ap}^{\text{R}}$ | (26) |
| pCJR01 | pBC SK::rRNA with unique *Not*I cloning site; $\text{Cm}^{\text{R}}$ | (26) |
| pBlueKan+cysM<sup>Pro</sup>-*lysR* | pBlueKan+cysM<sup>Pro</sup> with Gibson-cloned *lysR*; $\text{Kn}^{\text{R}}$; $\text{Ap}^{\text{R}}$ | This study |
| pCJR01comp-*lysR* | pCJR01 with cloned *aph*(3), *PcysM*, and *lysR*; $\text{Kn}^{\text{R}}$; $\text{Cm}^{\text{R}}$ | This study |

### *C. jejuni* motility assay

*C. jejuni* motility was measured by the ability of the bacterial strains to move through Brucella media plates containing a low agar concentration (0.3%). For strains possessing a cloned *aph*(3) gene, kanamycin (15 μg/mL) was added to the low agar Brucella plates. Low agar Brucella plates containing 5, 10 or 20 mM butyrate were made by diluting the appropriate amount of a 10 M, filter sterilized, solution of sodium butyrate (Sigma-Aldrich Inc. St. Louis, MO) into molten low concentration Brucella agars cooled to ∼60°C. The liquid Brucella agars containing butyrate were then poured into 100 mm round sterile plastic petri plates (Fisher Scientific, Waltham, MA) and allowed to cool overnight. Initially, individual bacterial strains were grown overnight in 3 mL of Brucella media. The next day 2 μL drops from the individual *C. jejuni* cultures were placed in the center of the appropriate 0.3% agar Brucella plates. The resulting plates were incubated for 16 h at 42°C in a microaerobic atmosphere chamber. Plates were then removed from the chamber and the diameters of the resulting circular bacterial migration zones across the plates were measured to the closest millimeter. Averages of individual strain measurements were calculated from three or more experimental replicates.

### *C. jejuni* attachment/biofilm assay

The ability of *C. jejuni* strains to attach to abiotic surfaces in the presence or absence of butyrate was measured using a version of the common crystal violet staining method (25). Strain CDC9511was grown overnight in Brucella broth. The next day, the overnight cultures were diluted 1:100 into fresh Brucella media containing 0, 5 mM or 20 mM butyrate. Seven 100 μL replicates of each sample were placed into individual wells of an untreated 96 well Nunc polystyrene plate (ThermoFisher Scientific, Waltham, MA) and the plate was covered with a rayon film (VWR, Radnor, PA). Negative control wells containing only 100 μl of Brucella broth were included on each plate. Plates were placed at 42°C in a microaerobic chamber for 96 h. After incubation, the supernatant of each well was carefully removed and the wells subsequently washed twice with 200 μLs of sterile water. Next, the remaining attached bacteria in each well were heat fixed at 60°C for 30 min. The wells and attached bacteria were then stained with 200 μLs of a 0.1% crystal violet solution for 30 min at room temperature. After which the stain was removed and the wells were again washed twice with 200 μL of sterile water. The remaining stain sequestered by the adherent bacteria in each well was then released by adding 200 μLs of 95% ethanol to each well. The remaining amounts of stain in each well was then determined by measuring the optical density (at 590 nm) of the liquid in each well using a Tecan Safire 2 plate reader (Männedorf, Switzerland). Average values were calculated from seven technical replicates from each of at least three experimental replicates to determine strain CDC9511’s ability to attach to the polystyrene well surfaces in the presence or absence of two different butyrate concentrations.

### Proteomic expression assay

The relative concentrations of the individual proteins, comprising the two different proteomes produced by incubating strain CDC9511 with or without exposure to 20 mM sodium butyrate, were determined using mass spectrometry. A 25 mL culture of strain CDC9511 was grown overnight microaerobically at 42°C. The resulting culture was separated into six 4 ml aliquots that were centrifuged at 7000 x g for 5 min to pellet cells. Three of the pellets were each resuspended in 6 mLs of fresh Brucella broth and the other three pellets were each resuspended in 6 mLs of Brucella broth plus 20 mM butyrate. This produced three experimental replicates for the two different conditions: with or without 20 mM butyrate. The samples were then incubated four hours at 42°C in a microaerobic chamber. After incubation the samples were centrifuged at 7000 x g to pellet cells, which were then washed twice in ProteaPrep (Protea Biosciences, Inc. Morgantown, WV) wash buffer (pH 8.0). ProteaPrep cell lysis buffer plus 100 μLs of a protease inhibitor cocktail was used to suspend each of the cell pellets before placing them in 2 mL micro-centrifuge tubes preloaded with low protein binding 100 micron zirconium beads (OPS Diagnostics, LLC, Lebanon, NJ). Next the cells were disrupted by agitation of the beads using a BeadBeater system (BioSpec Products, Bartlesville, OK) for 3 × 60 sec with at least 5 min rest periods on ice between treatments. The lysed samples were centrifuged at 13000 x g for 10 min at 4°C and the individual supernatants were collected. Supernatants were acidified to a pH of 2.5 using 10% v/v formic acid and incubated at room temperature for 10 min to inactivate the detergent present in the lysis buffer. Next, 50 mM ammonium bicarbonate was used to neutralize (pH 8.0) the protein samples. The protein concentration in the neutralized protein preps were measured using a BCA protein assay (Pierce, IL). The protein preps were reduced, alkylated and digested with trypsin (TrypsinGold, Promega,) for 12 h at 37°C following the manufacture’s protocols. The resulting tryptic peptide samples were acidified with 10% v/v formic acid and stored at -20°C for continued analysis by mass spectrometry.

The peptide preparations, digested with trypsin, were analyzed using a Nano-Acquity UHPLC (Waters Co., Milford, MA) running in the trap mode and equipped with a 20 mm x 180 μm, 5 μm Symetry C18 trap column (Waters) and a 200 mm x 75 μm, 1.8 μm HSS T3 (Waters) analytical column with a gradient of water:acetonitrile (0.1% formic acid) running with the composition from 95:5 to 50:50 with a total duration of 60 min and a flow rate of 450 nL/min. The nano-UHPLC flow was directed to a Q-Exactive Plus mass spectrometer (Thermo Scientific, San Jose, CA) using a nanospray Flex Ion source (Thermo Scientific) to obtain the peptides MSMS spectra for protein identification.

Mass spectra were analyzed by Proteome Discover 2.3 ((Thermo Fisher Scientific). All MS/MS samples were analyzed using Sequest, version IseNode in Proteome Discoverer. Sequest was set up to search *C. jejuni* from UniProtKB/TrEMBL-jejuni+81-176 database fasta file (1630 entries) using trypsin as the digesting enzyme. Sequest was searched with a fragment ion mass tolerance of 0.020 Da and a parent ion tolerance of 10.0 PPM. Carbamidomethyl of cysteine was specified in Sequest as a fixed modification. Oxidation of methionine and acetyl of the n-terminus were specified in Sequest as variable modifications.

Scaffold (version Scaffold_4.9.0, Proteome Software Inc., Portland, OR) was used to validate MS/MS based peptide and protein identifications. Peptide identifications were accepted if they could be established at greater than 95.0% probability by the Scaffold Local FDR algorithm. Protein identifications were accepted if they could be established at greater than 99.9% probability and contained at least 2 identified peptides. Protein probabilities were assigned by the Protein Prophet algorithm.

### Transcription analysis of *lysR*

RNA was collected from *C. jejuni* strain CDC9511 incubated for 4 h in Brucella broth supplemented with 0, 5 mM or 20 mM sodium butyrate using the Ambion Ribo-Pure Bacteria Kit (ThermoFisher Scientific). The RNA samples were subsequently DNase I treated. Synthesis of cDNA was performed using Invitrogen’s SuperScript III First-Strand Synthesis Kit (ThermoFisher Scientific) and an Applied Biosystems Veriti thermocycler (ThermoFisher Scientific) following manufacturer’s instructions.

Next, two primer sets for the *rpoA* housekeeping gene, serving as an internal control, and two primer sets to measure *lysR* gene transcription were used for the RT-PCR studies (*rpoA* primer set 1: aacatctgcttatacgccaacag[forward], aaaggatgggctagggtgattc[reverse]; *rpoA* primer set 2: cagtgcttggccttttgagatc[forward], cgcagttggagcatatcctatg[reverse]; *lysR* primer set 1: aaagtccaactcatacagccaac[forward], tccaagtctttcaaataaaatgccatc[reverse]; *lysR* primer set 2: cttacctacaccaaaagctttag[forward], cccctaagagcatactttcgtc[reverse]. RT-PCR was performed in a 96-well plate using an Applied Biosystems QuantStudio 6 RT-PCR system (ThermoFisher Scientific). Reactions of 50 μLs in total volume were used which contained: 25 μL of SYBR Green PCR Master Mix (Catalog #4309155; ThermoFisher Scientific), 1.25 μL of a forward and reverse primer at 10 μM each, 2 μL of cDNA, and 20.5 μL nuclease-free water (Ambion). The following amplification program was utilized: initial denaturation step at 50°C for 2 min, followed by 40 cycles of 95°C for 10 min, 95°C for 15 sec, and 60°C for 1 min. QuantStudio Real-Time PCR software, v. 1.3 was used to visualize PCR results. To control for genomic DNA contamination in the cDNA samples, a “no amplification control” (no reverse transcriptase control) and a “no-template control” containing all of the reverse transcriptase PCR reagents except for the RNA template were included for each reaction plate. To determine relative gene expression, the cycle threshold (CT) value of the internal control gene (*rpoA*) was subtracted from that of the *lysR* amplified samples to produce the ΔCT. The ΔCT of the treated samples minus the ΔCT of the untreated samples produced the ΔΔCT values. Finally the derivative of each ΔΔCT value was calculated (2^−Δfx^) to determine the degree differences between the treated and control groups.

### Construction of a constitutively expressed LysR strain

An additional copy of the *lysR* gene under constitutive control was placed at a secondary location on the chromosome of *C. jejuni* strain RM3194 using a previously described method (26). To produce the desired strain, we first cloned *lysR* into the assembly vector pBlueKan+cysM^Pro^ using a Gibson assembly strategy (Table 1). The *gene lysR* (from strain RM3194), and the plasmid pBlueKan+cysM^Pro^ were amplified using primers designed with 15 nucleotide extensions. The resulting PCR amplified fragments were assembled using NEBuilder HiFi DNA Assembly Master Mix; with the amplified *lysR* gene and the amplified pBlueKan+cysM^Pro^ joined to produce plasmid pBlueKan+cysM^Pro^-*lysR*. Next, the *lysR* expression cassette (kan gene, cysM promoter, *lysR* gene) was excised from plasmid pBlueKan+cysM^Pro^-*lysR* at the flanking *Not*I sites and transferred into the unique *Not*I site of pCJR01 producing the suicide integration plasmid pCJR01comp-*lysR*. The *lysR* gene was recombined into the rRNA spacer region of strain RM3194 by a double crossover integration event accomplished using a standard electroporation based transformation protocol (27). Proper integration of the expression cassette into the rRNA spacer region of the chromosome were confirmed by the presence of kanamycin resistance and the absence of chloramphenicol resistance in the transformed cells. The resulting strain was designated RM3194+*lysR*-express (Table 1). In order to better control for possible unintended side effects produced by the kanamycin gene and *cysM* gene promoter included with the *lysR* gene, in strain RM3194+lysR-express, we created strain RM3194+express by placing the same expression cassette previously used to make RM3194+*lysR*-express, only this time lacking the *lysR* gene, into wild-type RM3194. Experiments in this manuscript comparing the behavior of RM3194+*lysR*-express were therefore compared to RM3194+express instead of the wild-type RM3194 to better isolate the influence of the *lysR* gene on the strain.

Initially efforts had been made to inactivate the native copy of the *lysR* gene by homologous recombination using a construct that included sequence from either side of the *lysR* gene surrounding a gene coding for tetracycline resistance (*tetO*). Multiple attempts at this process failed to produce a successful double recombination/gene inactivation event. It is suggestive that the *lysR* gene may be essential in *C. jejuni*.

### Statistical analysis

All mean and standard deviation values reported in this paper were produced from at least three separate experimental replicates. Statistical analysis to compare experimentally related mean values were performed using multiple unpaired t-test comparisons with unequal variance. Significant differences were defined as P-values ≤ 0.05.

## Results

### Butyrate reduces *Campylobacter* motility

Two representative strains were selected after a review of the motility of *C. jejuni* strains in our laboratory collection (data not shown). One strain CDC9511 demonstrated motility in the upper third of our collection, ∼60 mm zones of migration after 16 h of incubation. The other strain CDC279’s motility was in the lower third of our collection, producing ∼30 mm zones of migration after 16 h incubation. The two stains with different baseline motility were utilized to determine the influence that butyrate has on *C. jejuni* motility.

Motility assays in low concentration Brucella agar plates with the addition of either 5 or 20 mM sodium butyrate produced significant reductions in the zones of migration compared to motility on plates without added butyrate (Figure 1). Additionally, the reductions in motility of both strains were also significantly different when comparing the zones of migration on the 5 mM butyrate plates versus the 20 mM plates. The overall magnitude in the butyrate mediated reductions in motility of CDC9511 appear much larger compared to that in strain CDC279. However, this is likely the result of strain CDC9511 having a significantly greater average baseline motility of 60.1 mm compared to CDC279’s baseline of 33.4 mm. On plates containing 20 mM butyrate the average motility of both strains are not that much different, 14.4 mm (CDC9511) and 12.8 mm (CDC279). Since the baseline motility of CDC9511 is almost double that of CDC279 the percentage of the reduction in motility is 76% for CDC9511 versus 62% for CDC279. The significant reductions recorded in the motility of strains CDC9511 and CDC279 caused by butyrate were repeated when we tested the motility of a larger set of *C. jejuni* strains from our laboratory collection during additional experimentation (data not shown).

**Figure 1:**
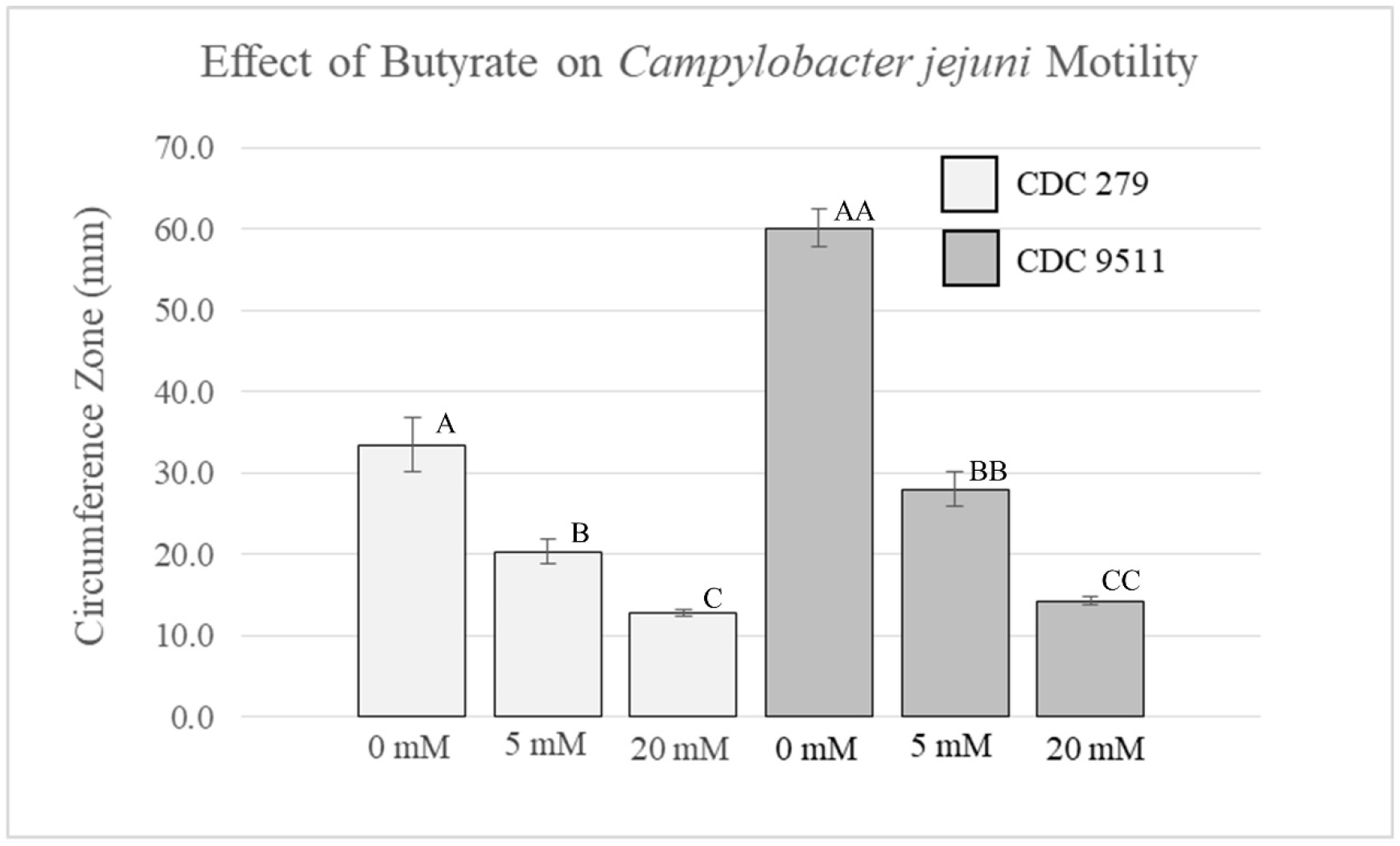
Average zones of motility in low concentration Brucella agar (0.3%) for *C. jejuni* strains CDC279 and CDC9511. Agar was supplemented with 0, 5 or 20 mM sodium butyrate. Significantly different values, in millimeters, are indicated by different letter notations.

To address the possibility that a growth defect in *C. jejuni* caused by butyrate was responsible for the decrease in observed motility, growth assays were performed for the strains in the presence of 5 or 20 mM butyrate. No significant effects, on the growth of strains CDC279 or CDC9511 in Brucella broth supplemented with sodium butyrate (5 or 20 mM), were observed as compared to the same strains grown in Brucella broth without butyrate (data not shown).

### Butyrate reduces *C. jejuni* attachment

The strain with the greater baseline motility, CDC9511, was used to determine if the two previous concentrations of butyrate had an effect on *C. jejuni’s* ability to attach to an abiotic surface. The ability of CDC9511 to attach to the walls of a polystyrene plate after 96 h was determined by measuring the optical density of crystal violet dye retained by the attached cells. The difference in the average absorbance of the retained dye from attached cells grown without butyrate (OD_590_ = 2.7) and the average from cells grown in 5 mM butyrate (OD_590_ = 2.4) was not large but it was significant (Figure 2). When the CDC9511 cells were grown in 20 mM butyrate the cells attaching to the well surface were significantly reduced by a much greater amount producing an average OD_590_ value of only 1.1. The results for the 20 mM butyrate incubated cells were also significantly different from both the no butyrate and 5 mM butyrate incubated cells (Figure 2). Similar experiments using incubation periods of 72 hours and less produced inconsistent results that suggested that butyrate was influencing the cells attaching to the well surfaces but were not significant upon analysis (data not shown). This result suggests that it might not just be attachment of the cells to the well surface that butyrate is affecting but also further development of the attached CDC9511 cells into a biofilm. As in the motility study, subsequent attachment studies using additional *C. jejuni* strains from our collection produced similar results with 20 mM butyrate reducing attachment or ensuing biofilm formation on the wells (data not shown)

**Figure 2:**
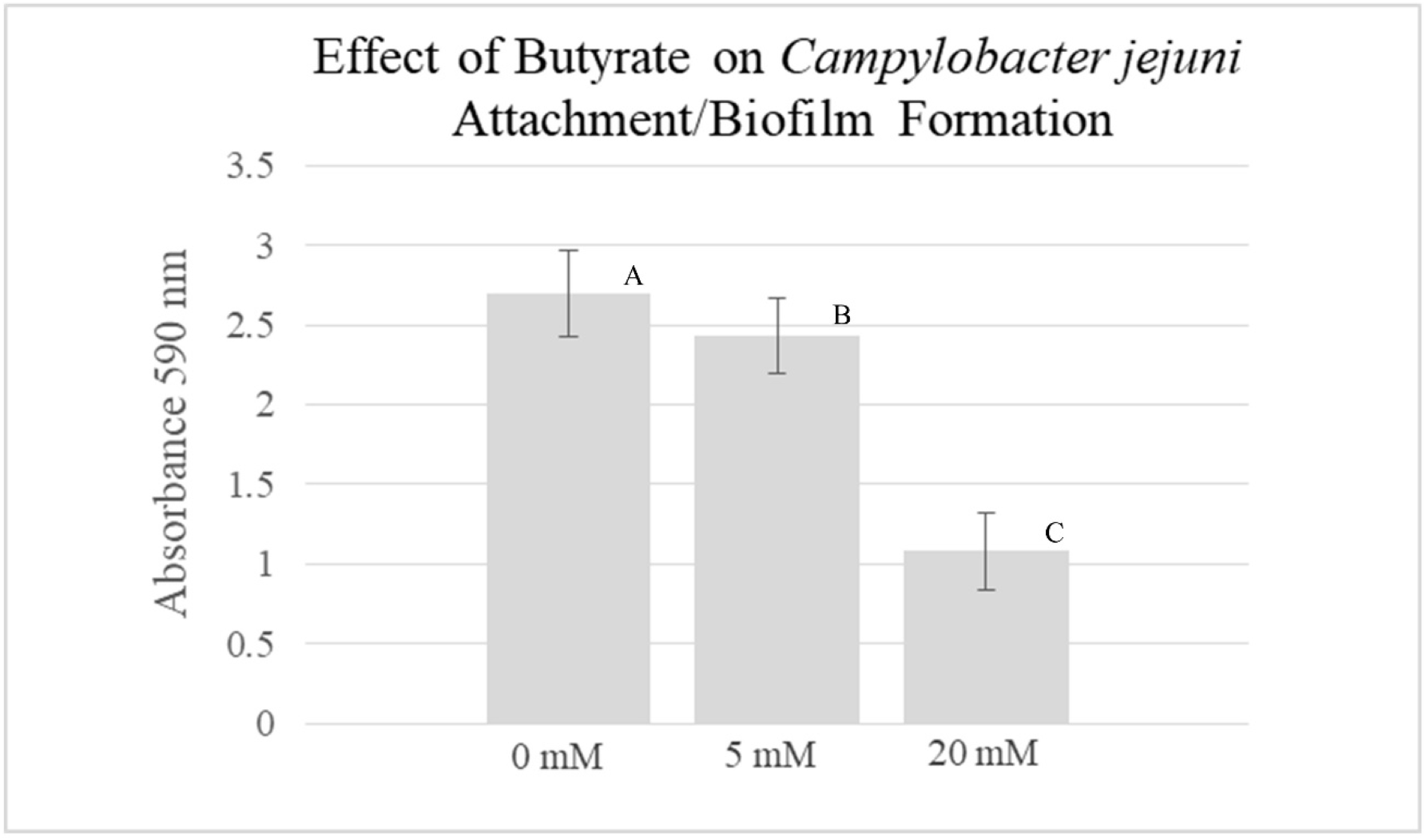
Average values for attachment to polystyrene well surface for strain CDC9511 exposed to 0, 5, or 20 mM butyrate. Significantly different OD_590_ values are indicated by different letter notations.

### Butyrate affects the expression of some CDC9511 proteins

Strain CDC9511 was utilized to compare the expression differences of individual proteins comprising the two unique proteomes produced by incubation of the strain in the presence or absence of 20 mM sodium butyrate. The expression differences of 799 individual proteins were compared in this study. *Campylobacter jejuni* strains possess ∼1600-1700 genes, therefore this study’s coverage of the potential proteome is approaching 50%. Comparisons of the protein expression levels from multiple replicates from the two different growth conditions determined that the expression of 31 proteins were significantly down regulated and 11 proteins were significantly up regulated in the presence of 20 mM sodium butyrate (Table 2).

**Table 2:** List of proteins upregulated or downregulated in the presence of 20 mM butyrate.

| Protein Accession # | Description | Molecular Weight | Spectrum ratio<br>(sample with<br>butyrate / sample<br>without butyrate) |
| --- | --- | --- | --- |
| **Upregulated in presence of 20 mM butyrate** |  |  |  |
| A0A0H3P9L4_CAMJJ | Primosomal protein<br>*priA* | 71 kDa | 2.0 |
| A0A0H3PBL5_CAMJJ | Shikimate kinase *aroK* | 19 kDa | 2.0 |
| A0A0H3PA26_CAMJJ | Peptidyl-arginine<br>deiminase family<br>protein | 38 kDa | 2.1 |
| A0A0H3PH57_CAMJJ | General glycosylation<br>pathway protein *pglH* | 41 kDa | 2.2 |
| A0A0H3PAB4_CAMJJ | Amidohydro-rel<br>domain-containing<br>protein | 31 kDa | 2.2 |
| A0A0H3PHT3_CAMJJ | Toluene tolerance<br>protein, putative | 22 kDa | 2.3 |
| A0A0H3PIZ5_CAMJJ | Uncharacterized<br>protein | 46 kDa | 3.0 |
| A0A0H3PAC7_CAMJJ | NADH-quinone<br>oxidoreductase, M<br>subunit *nuoM* | 56 kDa | 3.4 |
| A0A0H3PDC1_CAMJJ | RNA polymerase<br>sigma-54 factor *rpoN* | 49 kDa | 4.0 |
| A0A0H3PAB6_CAMJJ | Uncharacterized<br>protein | 13 kDa | 4.0 |
| A0A0H3PE64_CAMJJ | Membrane protein,<br>putative | 39 kDa | Spectra only<br>detected in +<br>butyrate sample |
| **Down regulated in presence of 20 mM butyrate** |  |  |  |
| A0A0H3PAM7_CAMJJ | Acetyltransferase,<br>GNAT family | 17 kDa | 0.1 |
| A0A0H3PDV7_CAMJJ | Selenocysteine-specific<br>elongation factor *selB* | 69 kDa | 0.1 |
| A0A0H3PCW1_CAMJJ | High affinity branched-<br>chain amino acid ABC<br>transporter, permease<br>protein *livM* | 38 kDa | 0.1 |
| A0A0H3PEQ4_CAMJJ | Uncharacterized<br>protein | 41 kDa | 0.2 |
| A0A0H3PAU6_CAMJJ | Dihydroorotase *pyrC* | 38 kDa | 0.2 |
| A0A0H3PIY1_CAMJJ | Uncharacterized | 10 kDa | 0.2 |

|  | protein |  |  |
| --- | --- | --- | --- |
| A0A0H3PIZ8_CAMJJ | Flagellar hook-basal body complex protein *fliE* | 11 kDa | 0.3 |
| A0A0H3PBD0_CAMJJ | Adenosylmethionine--8-amino-7-oxononanoate aminotransferase *bioA* | 49 kDa | 0.3 |
| Q2M5Q3_CAMJJ | Uncharacterized protein | 47 kDa | 0.3 |
| Q2TJD0_CAMJJ | LgtF | 61 kDa | 0.3 |
| A0A0H3PAB3_CAMJJ | Transcriptional regulator, LysR family | 34 kDa | 0.3 |
| A0A0H3PAT6_CAMJJ | Phosphatase, Ppx/GppA family | 37 kDa | 0.3 |
| A0A0H3PAZ9_CAMJJ | Thioredoxin, homolog | 18 kDa | 0.3 |
| A0A0H3PCD8_CAMJJ | Cation efflux family protein | 33 kDa | 0.3 |
| A0A0H3PAT2_CAMJJ | tRNA-dihydrouridine synthase *dusB* | 34 kDa | 0.3 |
| A0A0H3PIY4_CAMJJ | Shikimate dehydrogenase (NADP(+)) *aroE* | 30 kDa | 0.3 |
| A0A0H3PAA7_CAMJJ | Cytochrome d ubiquinol oxidase, subunit II *cydB* | 42 kDa | 0.3 |
| A0A0H3PAB1_CAMJJ | Membrane protein, putative | 25 kDa | 0.3 |
| A0A0H3PF42_CAMJJ | Penicillin-binding protein | 68 kDa | 0.4 |
| Q1G2W3_CAMJU | Type II restriction-modification enzyme *cju07* | 145 kDa | 0.4 |
| Q2A946_CAMJJ | Putative ATP/GTP binding protein | 21 kDa | 0.4 |
| A0A0H3PC40_CAMJJ | ATP-dependent DNA helicase RecG | 70 kDa | 0.4 |
| A0A0H3PDQ6_CAMJJ | AHS2 domain-containing protein | 36 kDa | 0.4 |
| A0A0H3P9C0_CAMJJ | ABC transporter, ATP-binding protein | 24 kDa | 0.4 |
| A0A0H3PAJ4_CAMJJ | Histidine biosynthesis bifunctional protein HisIE | 23 kDa | 0.4 |
| A0A0H3PBY2_CAMJJ | ThiF family protein | 24 kDa | 0.4 |
| Q2TJD2_CAMJJ | Lipopolysaccharide heptosyltransferase I<br>*waaC* | 39 kDa | 0.4 |
| A0A0H3PHE9_CAMJJ | *flgL*, flagellar hook-associated protein | 82 kDa | 0.5 |
| A0A0H3PAU4_CAMJJ | Molybdopterin<br>molybdenumtransferase | 43 kDa | 0.5 |
| A0A0H3PB85_CAMJJ | Transferase,<br>hexapeptide repeat<br>family | 20 kDa | 0.5 |
| A0A0H3PEA5_CAMJJ | Macrolide-specific<br>efflux protein macA | 43 kDa | 0.5 |

Among the down regulated proteins were two proteins: FliE, a flagellar hook basal body complex protein and FlgL, a flagellar hook associated protein. Both of these proteins are required for flagella mediated motility, and their apparent down regulation is therefore consistent with butyrate’s reductive effect on *C. jejuni* motility. Also of interest among the down regulated proteins is a protein belonging to the LysR family of transcription regulators. Given the wide range of additional genes a transcriptional regulator can influence, the down regulation of this protein could potentially have a far reaching influence on the *C. jejuni’s* response to butyrate exposure. There were also several currently uncharacterized proteins among those identified as being influenced by butyrate; with three of these uncharacterized being down regulated and two of them being up regulated.

### Butyrate does not negatively impact l*ysR* transcription

In an effort to determine if the previously observed butyrate mediated down regulation of the LysR protein occurred at the level of transcription we measured the *lysR* transcription levels of CDC9511 with or in the absence of 5 or 20 mM sodium butyrate using RT-PCR. The CDC9511 cells used for the RT-PCR assays were incubated in the same manner as the cells used for the proteomic comparison with exposure times to the butyrate always being 4 h. Transcription studies produced results contradictory to those expected if the previously observed down regulation of the LysR protein occurred at a transcriptional level. The results of the studies showed that *lysR* transcription levels were increased in the presence of 20 mM butyrate (Table 3). Incubation in 20 mM butyrate produced 2.8 to 4.6 fold increases in *lysR* transcription which was somewhat dependent on the combination of *lysR* and *rpoA* primer sets. Additionally, the changes in transcription levels were very inconsistent at the 5 mM butyrate challenge level, with the results depending greatly on the combination of *lysR* and *rpoA* control primer pairs.

**Table 3:**
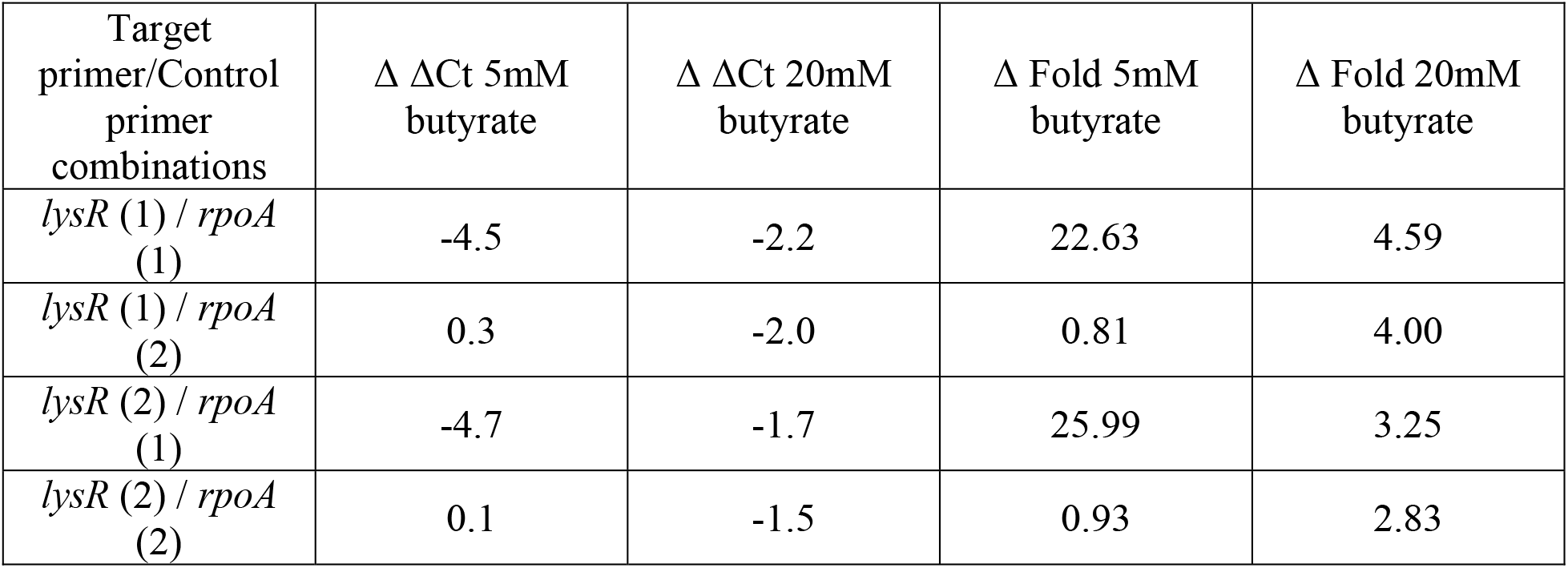
Change in transcription levels (by RT-PCR) of *lysR* exposed to 5 or 20 mM butyrate.

### Constitutive *lysR* expression increased motility in the presence of butyrate

A copy of the *lysR* gene was placed under the control of a constitutive promoter expression cassette (Kan^R^-*cysM* promoter-*lysR*) within the rRNA spacer region of the chromosome of *C. jejuni* RM3194 and designated RM3194+*lysR-*express. An additional RM3194 had just the expression cassette portion (Kan^R^-*cysM* promoter) without the *lysR* gene also placed into its chromosome within the rRNA spacer region and designated RM3194+express. Therefore, comparisons between these two strains should only produce differences that are related to constitutive expression of the *lysR* gene. Both stain RM3194+*lysR-*express and RM3194+express were the subject of motility assays on low concentration Brucella + Kanamycin (15μg/ml) agar both with and without 10 mM Butyrate. Zones of migration for each strain in agar without butyrate were compared to zones of migration for the same strain in agar with 10 mM butyrate. For both strains the presence of butyrate reduced the motility of the bacteria compared to the strains baseline motility in low concentration agar without butyrate. However, the RM3194+*lysR-*express strain constitutively expressing *lysR* did not see as large of a reduction in motility compared to the RM3194+express strain not constitutively expressing *lysR* (Figure 3). The differences in motility in the presence of butyrate observed between the two strains was modest but significant. Strain RM3194+*lysR-*express on low agar plates containing 10 mM butyrate produced 75.0% of the motility it was able to produce on low agar plates without butyrate (baseline motility). While strain RM3194+express managed only 68.5% motility under the same conditions. Since the constitutive expression of *lysR* did not fully restore motility to the baseline levels present without butyrate it is unlikely that reduction of LysR expression is the only way in which butyrate is influencing *C. jejuni* motility. Additionally, the constitutive *lysR* expression may also be in part downregulated by some post-transcriptional system, limiting the production of functional LysR protein and its subsequent influence on motility.

**Figure 3:**
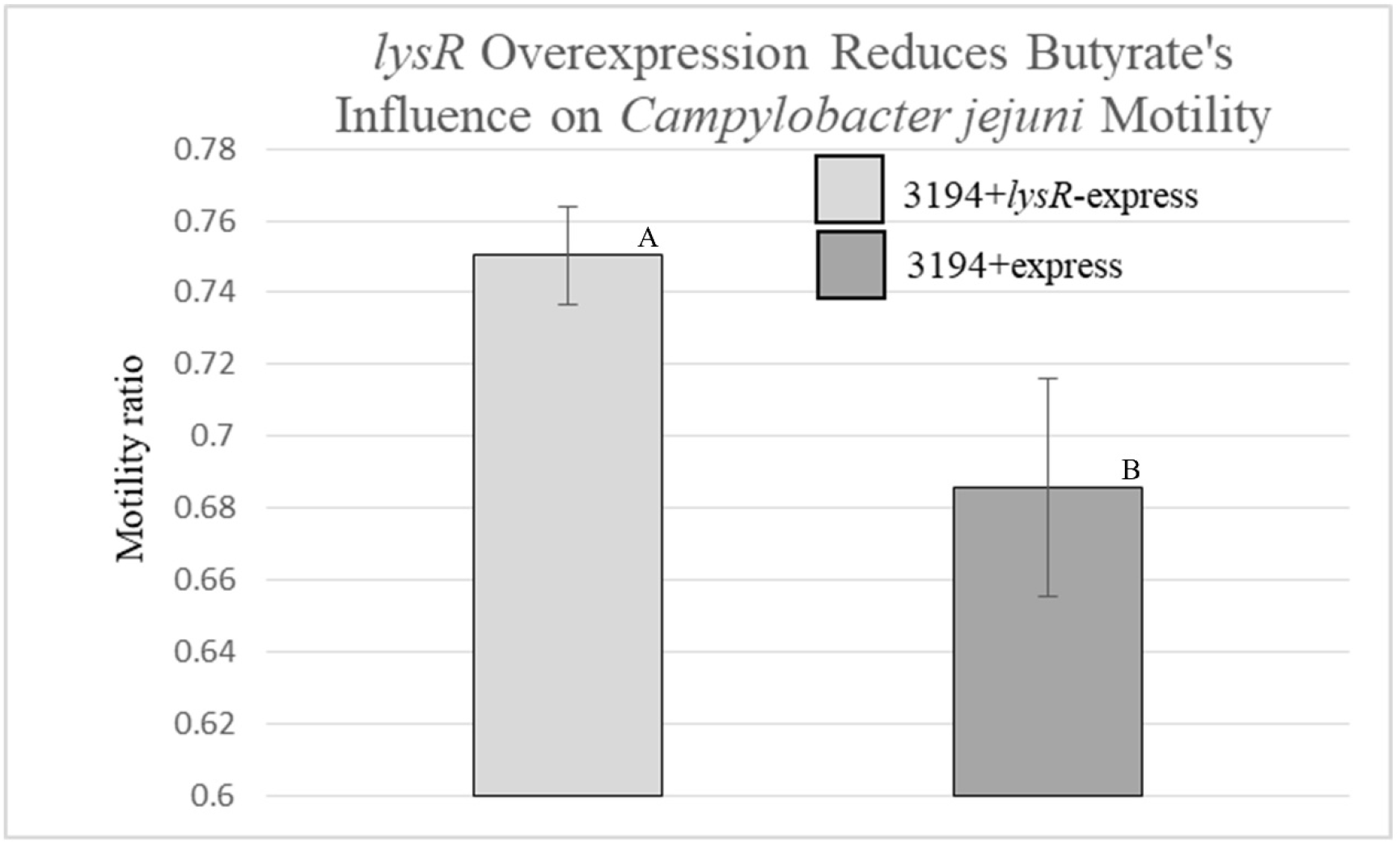
Recovery of motility in RM3194+*lysR*-express strain with an extra copy of the *lysR* gene with a constitutive promoter compared to RM3194+express, the same strain without the extra *lysR* copy. The average reported values are a ratio of the zone of motility in the presence of 10 mM butyrate divided by the zone of motility without butyrate. Significantly different ratio values are indicated by different letter notations.

## Discussion

The presented research suggest that butyrate negatively influences *C. jejuni’s* ability to move and adhere to surfaces. If true, this would definitely lend support to the view that butyrate is an inhibiter of *C. jejuni* mediated colonization and infection of the digestive tracts of chickens and humans. The reported negative influence of butyrate on motility and attachment was observed in the presence of both 5 and 20 mM concentrations of the SCFA. These concentrations of butyrate were selected because they were well within the physiological levels observed for butyrate in both chickens and humans. Concentrations of butyrate in the lower intestines of chickens have been observed to fall in the range of 20-30 mM (15). While in the human lower intestines, the combined concentrations of the three most prominent SCFAs, acetate, butyrate, and propionate, have been measured in the range of 70-140 mM (12, 28). The use of physiologically relevant levels of butyrate should reduce concerns that the reported results are an artifact resulting from levels of butyrate that *C. jejuni* would never be expected to encounter within its host environments.

Since *C. jejuni* flagella are needed both for motility and adhesion it was not surprising given butyrate’s influence on motility that it would also negatively impact surface adhesion. Flagella is a primary virulence factor for *Campylobacter’s* infection of the human intestinal tract (29). It is necessary for the process of reaching, adhering to and invading the epithelial cells of the intestines (30). While results were only provided for butyrate decreasing *C. jejuni* binding to an abiotic surface, it is reasonable to think that butyrate would also reduce binding to biotic surfaces given the importance of flagella in adherence. Research exists that looked at the influence of butyrate on *C. jejuni’s* invasion of cell monolayers but this research focused on the effects on invasion and translocation and not specifically attachment (17). Therefore, a study of butyrate and *C. jejuni* attachment to human cell monolayers may be a valuable research pursuit.

A potentially contradictory set of results presented in this paper are the proteomic expression results for the protein LysR and the transcriptional analysis of its corresponding gene *lysR* which codes for the protein. The LysR protein was first observed as being down regulated in the presence of butyrate; however when the transcription levels of the *lysR* gene were measured under the same conditions it appeared to be upregulated by butyrate. This is not overly concerning given that there is often a lack of correlation between proteomic and genomic expression data given the common existence of post-transcriptional regulatory systems in bacteria. As an example, activation of the *E. coli* BarA sensor by SCFA leads to the sequestering of the CsrA protein which normally acts to block translation of mRNAs, an illustration of the same sort of post-transcriptional regulation which may be happening with LysR (19). Of additional note, *C. jejuni* possesses a CsrA orthologue that negatively influences the expression of flagella if not bound by another protein, also a post-transcriptional example of regulation (30). It is therefore possible to imagine the recently described *C. jejuni* butyrate sensor responding to the presence of butyrate by activating the CsrA protein to tie up *lysR* gene transcripts and decrease the amounts of the LysR proteins, as we observed in this study. A decrease in LysR, a transcriptional factor, could then lead to a decrease in proteins essential for motility, such as the flagella FlgL and FliE proteins which are two proteins we also observed to be down regulated in this study. Unfortunately our inability to inactivate the *lysR* gene in *C. jejuni* makes it difficult to establish a definitive connection between LysR and flagellar gene expression. However, our observation of constitutive expression of *lysR* partially compensating for butyrate mediated reduction in motility suggests a relationship between LysR production and flagellar expression.

There is very strong evidence for the SCFA, butyrate, playing a positive role in *Campylobacter’s* colonization of the poultry digestive tract and infection of the human digestive tract (13, 14). However, there remains examples in the research where SCFA’s, including butyrate, have a negative influence on *C. jejuni* that could lead to difficulties in the process of colonization or infection (15-17). The research presented here also suggests that butyrate at physiologically relevant concentrations could have a negative influence of *C. jejuni* through the reduction of flagella activity. Additionally, the LysR protein appears to play a role in the regulation of flagella given that a strain with an additional constitutively promoted *lysR* gene regained some of its motility in the presence of butyrate. Future research should focus on determining if butyrate’s apparent negative influence on motility and attachment are outweighed by butyrate’s positive influence on other colonization and virulence factors. Additionally, it would be ideal if studies investigating the influence of SCFAs on *C. jejuni* could be extended to multiple strains isolated from both environmental and clinical sources that have been minimally passaged within laboratories. *Campylobacter jejuni* has a small fairly plastic genome capable of accumulating significant genomic variation. It is possible therefore, that not all strains respond to SCFAs in the same manner. Addressing this possibility may help clarify contradictory observations.

## Acknowledgments

This research is supported by congressional appropriated funds assigned to the USDA - Agricultural Research Service, National Program 108 Food Safety [Project # 8072-42000-094-000-D].

